# Benchmarking phasing software with a whole-genome sequenced cattle pedigree

**DOI:** 10.1101/2021.10.27.466052

**Authors:** Claire Oget-Ebrad, Naveen Kumar Kadri, Gabriel Costa Monteiro Moreira, Latifa Karim, Wouter Coppieters, Michel Georges, Tom Druet

**Affiliations:** Unit of Animal Genomics, GIGA-R and Faculty of Veterinary Medicine, University of Liège (B34), 4000 Liège, Belgium; Animal Genomics, ETH Zürich, Zürich, Switzerland; Genomics Platform, GIGA, University of Liège (B34), 4000 Liège, Belgium

**Keywords:** Haplotype, phasing, sequencing data, cattle

## Abstract

**Background:** Accurate haplotype reconstruction is required in many applications in quantitative and population genomics. Different phasing methods are available but their accuracy must be evaluated for samples with different properties (population structure, marker density, etc.). We herein took advantage of whole-genome sequence data available for a Holstein cattle pedigree containing 264 individuals, including 98 trios, to evaluate several population-based phasing methods. This data represents a typical example of a livestock population, with low effective population size, high levels of relatedness and long-range linkage disequilibrium.

**Results:** After stringent filtering of our sequence data, we evaluated several population-based phasing programs including one or more versions of AlphaPhase, ShapeIT, Beagle, Eagle and FImpute. To that end we used 98 individuals having both parents sequenced for validation. Their haplotypes reconstructed based on Mendelian segregation rules were considered the gold standard to assess the performance of population-based methods in two scenarios. In the first one, only these 98 individuals were phased, while in the second one, all the 264 sequenced individuals were phased simultaneously, ignoring the pedigree relationships. We assessed phasing accuracy based on switch error counts (SEC) and rates (SER), lengths of correctly phased haplotypes and pairwise SNP phasing accuracies (the probability that a pair of SNPs is correctly phased as a function of their distance). For most evaluated metrics or scenarios, the best software was either ShapeIT4.1 or Beagle5.2, both methods resulting in particularly high phasing accuracies. For instance, ShapeIT4.1 achieved a median SEC of 50 per individual and a mean haplotype block length of 24.1 Mb in the second scenario. These statistics are remarkable since the methods were evaluated with a map of 8,400,000 SNPs, and this corresponds to only one switch error every 40,000 phased informative markers. When more relatives were included in the data, FImpute3.0 reconstructed extremely long segments without errors.

**Conclusions:** We report extremely high phasing accuracies in a typical livestock sample of 100 sequenced individuals. ShapeIT4.1 and Beagle5.2 proved to be the most accurate, particularly for phasing long segments. Nevertheless, most tools achieved high accuracy at short distances and would be suitable for applications requiring only local haplotypes.

## Background

Haplotype phasing consists in the reconstruction of haplotypes inherited from each parent. On autosomes, diploid individuals carry two alleles (eventually identical) at polymorphic sites, each allele being inherited from one of the two parents. The combination of alleles from one of the homologous chromosomes is called haplotype, or phase. However, genotyping data obtained with genotyping arrays or from whole-genome sequencing experiments are typically unphased, the origin of each allele remaining unknown. Therefore, statistical phasing methods must be used to determine the set of alleles belonging to each homolog, that were co-inherited. Haplotype information can be used in many applications in quantitative and population genomics, including missing genotype imputation [1, 2], identification of identical-by-descent (IBD) segments in outbred or experimental populations [3–5], quantitative-trait *locus* (QTL) mapping [6], haplotype-based association studies [7–9] or genomic predictions [10–12], demographic inference [13, 14], identification of signatures of selection [15, 16], allele’s age estimation [17], estimation of linkage disequilibrium (LD) -based recombination maps [18] or identification of cross-over (CO) events in genotyped pedigrees [19, 20]. Haplotypes present indeed higher LD with underlying causative variants and allow the estimation of the length of shared IBD segments, a measure related to the number of generations to their common ancestor [21]. Haplotypic information is also required to understand interactions among tightly linked *loci*, for instance to study allele specific expression, identify deleterious compound heterozygotes or to determine how combinations of variants affect gene expression [22, 23].

Haplotype phasing methods have been reviewed by Browning and Browning [24]. They can be divided into two main groups: those relying on pedigree relationships (e.g., [25, 26]) and those than can be applied in samples of unrelated individuals by exploiting LD information, often referred to as population-based methods (e.g., [7,27,28]). Nevertheless, some methods present hybrid properties by exploiting both sources of information (e.g., [29–31]). Other methods apply heuristic rules by matching target haplotypes to libraries of reference haplotypes in windows (e.g., [30]). Long-range phasing methods use such heuristic approaches and rely also on the identification of surrogate parents [29, 32]. Most recent advances in phasing methods were related to their ability to handle huge data sets, including thousands of sequenced samples [33–35]. Phasing accuracy of these different approaches will impact the outcome of different haplotype-based applications and must be assessed. Although this is most often initially tested in human populations, it should ideally be realized in populations with different demographic histories and levels of relatedness.

Here we take advantage of a unique sequenced cattle pedigree to assess accuracy of several population-based phasing methods, including recent methods commonly used in livestock species. This sample is a typical example of a livestock population with reduced effective population size, high levels of relatedness and long-range LD, and containing 100 to 200 sequenced individuals. We show that such data can be phased with extremely high accuracy. In addition, we illustrate that the raw sequence data requires stringent filtering to obtain accurate haplotypes.

## Results

### Quality of whole-genome sequence genotype data after applications of different procedures

We assessed the quality of the genotyping data by comparing the number of identified CO with the evaluated data to the expected number of CO, obtained with a high-quality reference map validated with 115,967 genotyped individuals and 30,331 SNPs (see *Methods*). In both cases, CO were identified using a pedigree-based approach [36]. The ARS-UCD1.2 bovine genome assembly [37] was used as starting point for the reference map. A total of 18 SNPs showing evidence of incorrect map positions were then discarded. Among those, 13 matched with regions also flagged by Quanbari and Wittenburg [38] as potential errors in the genome build. After removal of these SNPs, we found no more evidence for map errors. Using this reference map and our sequenced pedigree containing 264 individuals, we found an average of 24 CO per individual, 26 and 23 in males and females, respectively.

When CO were estimated using the 15,327,429 SNPs that passed the variant quality score recalibration (VQSR) procedure (with the threshold set to 99.9), the average counts per meiosis were highly inflated, equal to 1,416 (**Figure 1**). When the threshold for the VQSR filtering was set to 97.5, resulting in the selection of 11,030,905 SNPs, the average number of CO dropped to 254 confirming that the data quality was improved. However, this value was 10 times larger than the expected values, clearly indicating that the sequence genotype data required further cleaning. Subsequent selection of a subset of 8,435,899 variants behaving like true Mendelian variants (see *Methods*), with genotype frequencies close to Hardy-Weinberg proportions, and with minor allele frequency higher than 0.01, resulted in the identification of 167 CO on average per meiosis, still six times above expectations. Refining genotype calls using Beagle4.1 [39] clearly improved the genotype quality, the number of identified CO being reduced by a factor 3 (51 CO per meiosis on average). We then removed regions presenting high coverage, excessive levels of recombination or of genotyping errors (see *Methods*), resulting in the removal of 18,220 additional SNPs. The total number of SNPs was then 8,417,679 ranging from 166,292 (chromosome 25) to 529,626 (chromosome 1) per chromosome (**Table S1** in **Additional file 1**). This removal further improved the quality of our data as the number of identified CO dropped to 31 CO per meiosis on average. Finally, by setting remaining genotypes that were discordant in parent-offspring pairs to missing (e.g., opposite homozygous), the average number of identified CO was further reduced to 30 CO. Overall, the application of these different procedures allowed us to reduce the average number of CO from 1,416 to 30 CO, closer to expectations. Nevertheless, we still identified on average 5 additional CO with the final sequence data compared to results obtained with the high-quality reference map. These 5 additional CO might be missed with the 50K map but could also correspond to errors, meaning that haplotypes obtained with a family-based approach might still contain a few errors.

**Figure 1.**
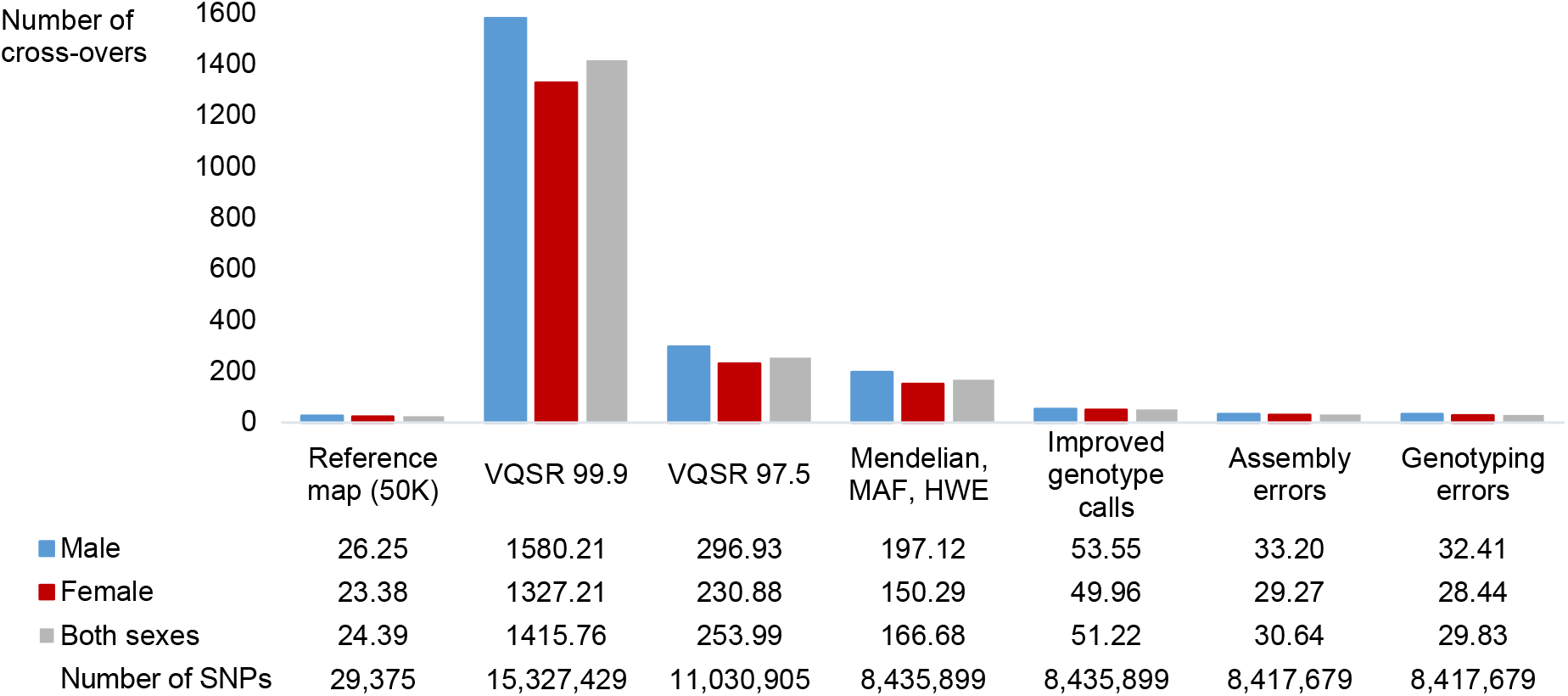
Quality of whole-genome sequence genotype data after applications of different procedures. Average cross-over counts per meiosis obtained with a high-quality reference map (50K) and after application of different procedures to improve the quality of the whole-genome sequence SNPs (see Methods).

### Comparison of phasing quality achieved with different population-based phasing methods

#### Strategy

To assess the phasing quality from different LD-based methods we used the haplotypes from 98 sequenced individuals that had both their parents also sequenced (sequenced trios). The haplotypes of these sequenced offspring (validation individuals) were phased using Mendelian rules, that are exact in absence of genotyping errors, to serve as the “true haplotypes”. Population-based approaches were then applied in two scenarios, either with only the 98 validation individuals (scenario 1), or with the full data set consisting in 264 individuals, but ignoring the pedigree relationships (scenario 2). Most phasing metrics are computed with respect to heterozygous markers phased in the true haplotypes (the gold standard) with Mendelian rules since these markers are informative. On average, each of the 98 individuals had 1,964,220 such informative markers in both scenarios (**Table S1** in **Additional file 1**). The different metrics used to assess phasing quality are described in the *Methods* section and most of them are illustrated in **Figure 2**.

**Figure 2.**
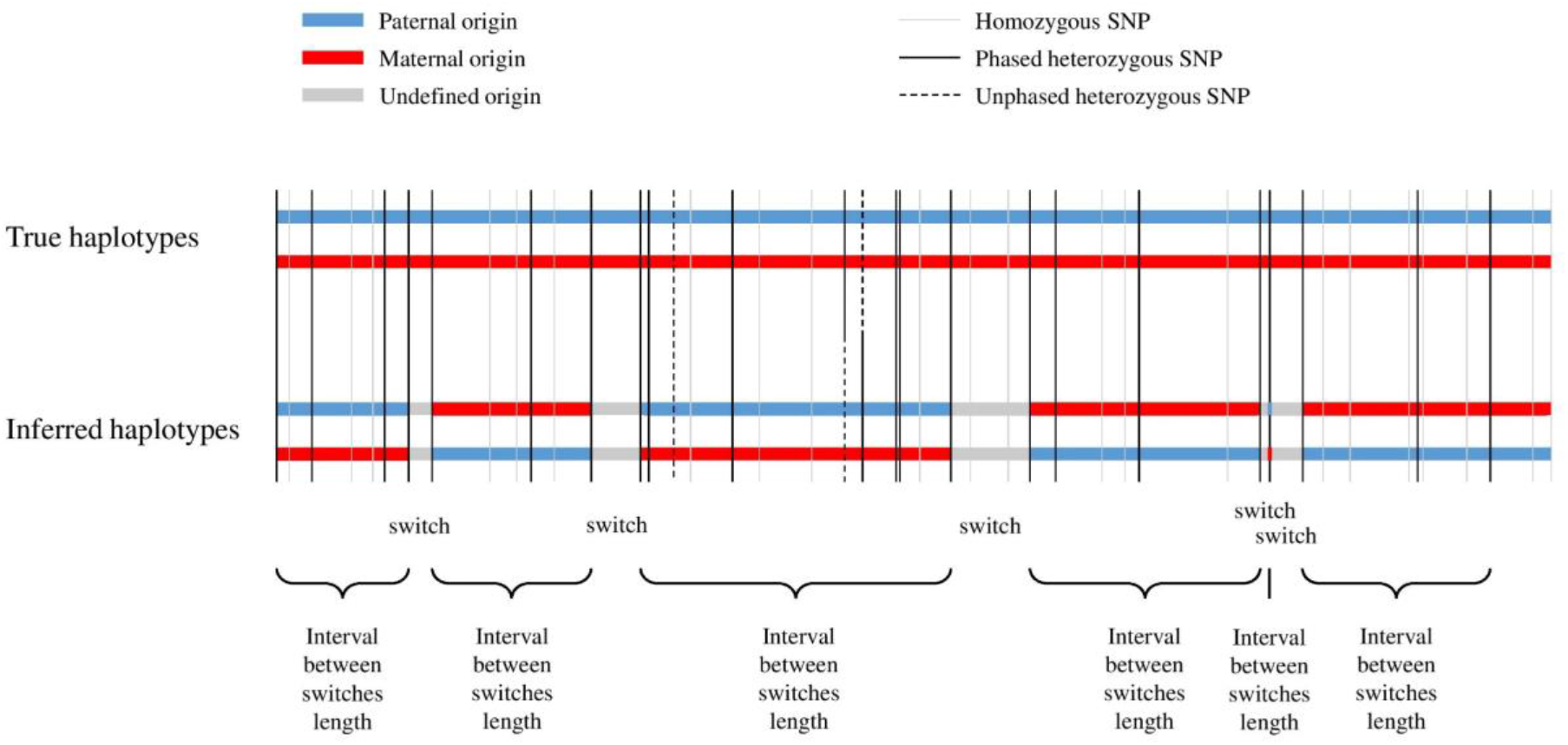
Illustration of different metrics used to assess phasing quality. Illustration of the switches between true haplotypes (gold standard obtained using Mendelian rules) and inferred haplotypes (estimated by the different population-based phasing methods). The quality adjusted (QA) haplotype block length computed in this study is equal to the product between the distance between switches and the phasing yield (proportion of informative SNPs that are phased in inferred haplotypes, the informative SNPs being the SNPs that are heterozygous and phased in the true haplotypes).

#### Phasing yield

Most of the tested phasing methods achieved 100% phasing yield, and this was almost true also for FImpute3.0 that phased on average more than 99.99% of the heterozygous SNPs in both tested scenarios. Only AlphaPhase1.3 failed to phase all the SNPs, with 1.31% remaining unphased in the first scenario for instance.

#### Switch error count (SEC) and rate (SER)

Median switch error count (SEC) and rate (SER) computed on the 98 validation individuals are provided for the main phasing algorithms and each scenario in **Figure 3** and **Table 1**. When only the 98 validation individuals were used for phasing, the median SEC was around 4,500 to 5,000 for a group of methods including AlphaPhase1.3, Eagle2.4 and FImpute3.0. These values correspond to a SER slightly below 0.25%, meaning that switch errors occur on average every 400 informative markers. Haplotypes obtained with Beagle4.1 had clearly lower SEC than these first methods, close to 2,750. ShapeIT4.1 performed even better with median SEC and SER below 400 and 0.02% respectively, corresponding to one switch error every 5,000 informative markers. Finally, Beagle5.2, relying on a new algorithm compared to earlier Beagle versions (until Beagle4.1), resulted in the lowest SEC and SER, outperforming all other methods, the median values were below 200 and 0.01% per individual, corresponding to one switch every 10,000 informative markers. This represents a reduction by a factor 20 compared to AlphaPhase1.3, Eagle2.4 and FImpute3.0, whereas ShapeIT4.1 generated ten times fewer switches than these methods. When methods are compared with other summary statistics related to SEC and SER such as the mean, minimal and maximal values or as their range (**Figure 3A** and **Tables S2-S3** in **Additional file 1**), the ranking of the method remained similar with Beagle5.2 performing best, followed by ShapeIT4.1. When comparing different versions of the same software in terms of SEC and SER (**Figure S1A** in **Additional file 1**), we observed that newer versions are more accurate as expected. In particular, AlphaPhase1.3 represents a major improvement with respect to AlphaPhase1.1. Regarding Beagle’s versions relying on a directed acyclic graph, Beagle3.3 and Beagle4.0 had close performances and Beagle4.1 appeared as an important improvement. Several of these versions of Beagle presented a lot of variation among individuals. Regarding the latest Beagle’s versions, relying on the Li and Stephens model [40], Beagle5.1 was slightly better than Beagle5.0 whereas Beagle5.2 represented a substantial improvement. Finally, ShapeIT2 and ShapeIT4.1 achieved similar performances.

**Figure 3.**
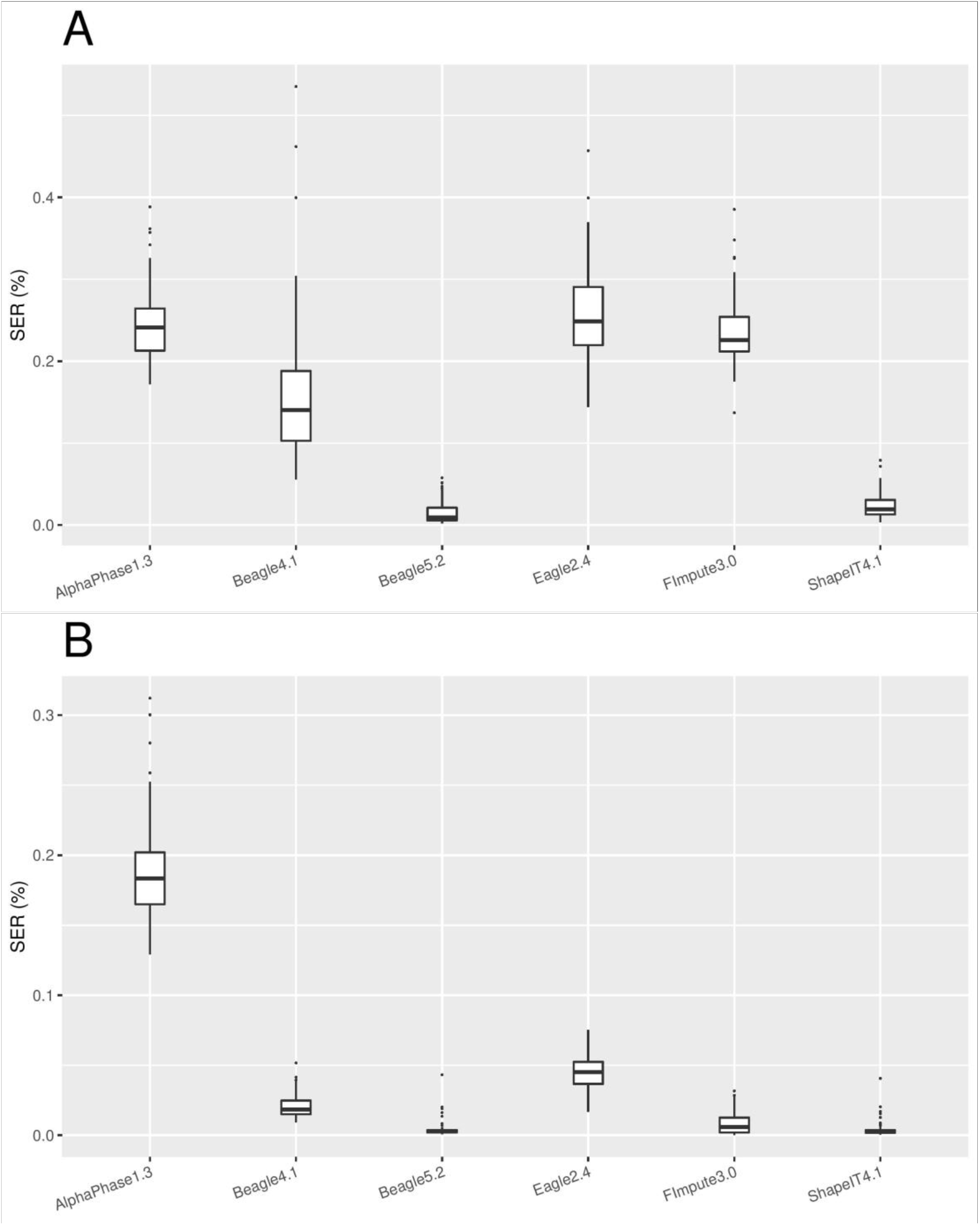
Switch error rates in both scenarios. Boxplots of switch error rates (SER, %) obtained with AlphaPhase1.3, Beagle4.1, Beagle5.2, Eagle2.4, FImpute3.0 and ShapeIT4.1, computed for the 98 validation individuals (A) in scenario 1 with only the 98 validation individuals, and (B) in scenario 2 with the 264 sequenced individuals.

**Table 1.**
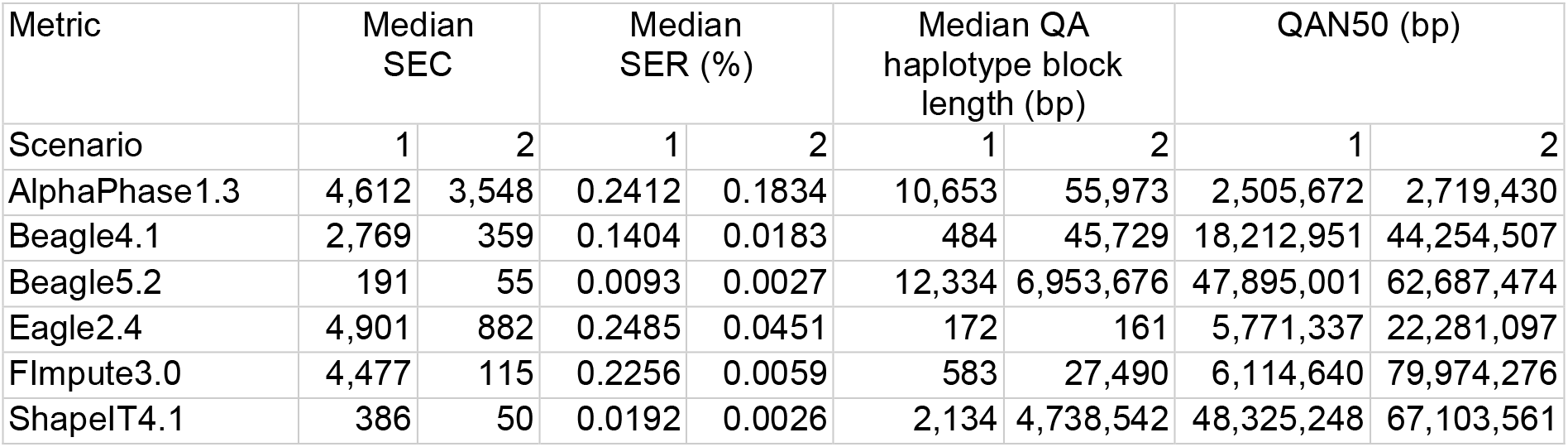
Results of different metrics used to assess phasing quality in both scenarios. Median values of switch error counts (SEC), switch error rates (SER, %), quality adjusted (QA) haplotype block length (bp), and QAN50 (bp), obtained with AlphaPhase1.3, Beagle4.1, Beagle5.2, Eagle2.4, FImpute3.0 and ShapeIT4.1, computed for the 98 validation individuals in each scenario (scenario 1: using the 98 validation individuals; scenario 2: using the 264 sequenced individuals).

When a larger data set was used for phasing, consisting in 264 individuals including the sequenced parents (scenario 2), the phasing accuracy improved for most of the methods (**Figure 3B** and **Table 1**), with less variation among individuals. For AlphaPhase1.3, the SEC reduction remained however modest and it consequently ranked last. For all the other phasing methods, the median SEC was below 1,000, around 900 and 350 for Eagle2.4 and Beagle4.1, respectively. FImpute3.0 showed the highest improvement compared to scenario 1, the median SEC being reduced by almost 40 folds and dropping to 115. However, Beagle5.2 and ShapeIT4.1 still performed best with median SEC values equal to 55 and 50, respectively. These values correspond to extremely low median SER, equal to 0.0027 and 0.0026%, respectively, and to one switch error every 40,000 informative SNPs. The ranking remains similar with other summary statistics (**Figure 3B** and **Tables S2-S3** in **Additional file 1**), except that FImpute3.0 presented the lowest minimum and maximum individual SEC. With FImpute3.0, the lowest value was equal to 2, indicating that almost all chromosomes were perfectly phased for that individual. With ShapeIT4.1 and Beagle5.2, the best phased individual had only 12 and 13 SEC, respectively. The median SEC dropped for all the different versions of the tested software, and the variation among individuals was strongly reduced, in particular for Beagle3.3, Beagle4.0, Beagle4.1 and Beagle 5.1 (**Figure S1B** in **Additional file 1**). The ranking of these different versions, in terms of SEC or SER, was similar to the ranking observed in the first scenario, with the exception of ShapeIT2 that presented now higher SEC and variation levels than ShapeIT4.1. Differences between Beagle5.1 and Beagle5.2 were also smaller than in the first scenario.

#### Length of correctly phased haplotype blocks

Statistics relying on SEC does not provide a full description of their distribution along the chromosomes and of the resulting distribution of length of correctly phased haplotype segments. Therefore, we also computed the quality adjusted (QA) haplotype block length and the QAN50 metrics, as described in the *Methods* section, in order to highlight the ability of a phasing tool to produce long correctly phased blocks within a chromosome, without switch error.

In the first scenario, median QA haplotype block length are equal respectively to 10 and 12 kb with AlphaPhase1.3 and Beagle5.2, and clearly lower with other methods (**Table 1**). The ranking of the methods based on the median QA haplotype block lengths is thus very different from comparisons based on SEC, with AlphaPhase1.3 ranking second. However, when mean values are used in comparisons (**Table S4** in **Additional file 1**), the ranking follows results obtained with SEC. These mean lengths of correctly phased segments range from 500 kb with AlphaPhase1.3 to 7.5 Mb with Beagle5.2. Compared to AlphaPhase1.3, the median values were five times lower with ShapeIT4.1 but the haplotype block lengths were on average ten times longer. This indicates that some methods such as ShapeIT4.1 tend to produce a lot of small correctly phased segments (switch errors being close) in combination with very long correctly phased segments (up to 156.8 Mb with ShapeIT4.1, a full chromosome), whereas other methods such as AlphaPhase1.3 tend to provide more uniform distances between successive switch errors. The QAN50 metrics obtained for different methods (**Figure 4A** and **Table 1**) indicate that with AlphaPhase1.3, 50% of the genome is included in correctly phased segments longer than 2.5 Mb. The QAN50 increases to 5.8 and 6.1 Mb with FImpute3.0 and Eagle2.4, respectively (approximately 20,000 SNPs, **Figure 4C**) and to 18.2 Mb with Beagle4.1(approximately 60,000 SNPs). ShapeIT4.1 and Beagle5.2 performed best with a QAN50 close to 48 Mb corresponding to blocks of approximately 170,000 SNPs. **Figure 4A** provides the full distribution of QAN values (from 100 to 0%), with very similar curves for ShapeIT4.1 and Beagle5.2. It allows also to determine the percentage of the genome included in correctly phased segments longer than different thresholds as reported in **Table 2**. With Beagle5.2 for instance, 96.3, 93.6, 89.3 and 48.1% of the genome is included in correctly phased segments of at least 1, 5, 10 and 50 Mb, respectively. These values are close with ShapeIT4.1 and lower with the remaining methods, only 86.4, 54.0, 33.2 and 1.3% with Eagle2.4, for instance. Most recent versions of tested software performed better in terms of QA and QAN50 than older versions (**Figure S2A,C** and **Table S4** in **Additional file 1**), with the exception of ShapeIT2 that had similar statistics as ShapeIT4.1, and Beagle4.1 that presented better results than Beagle5.0.

**Figure 4.**
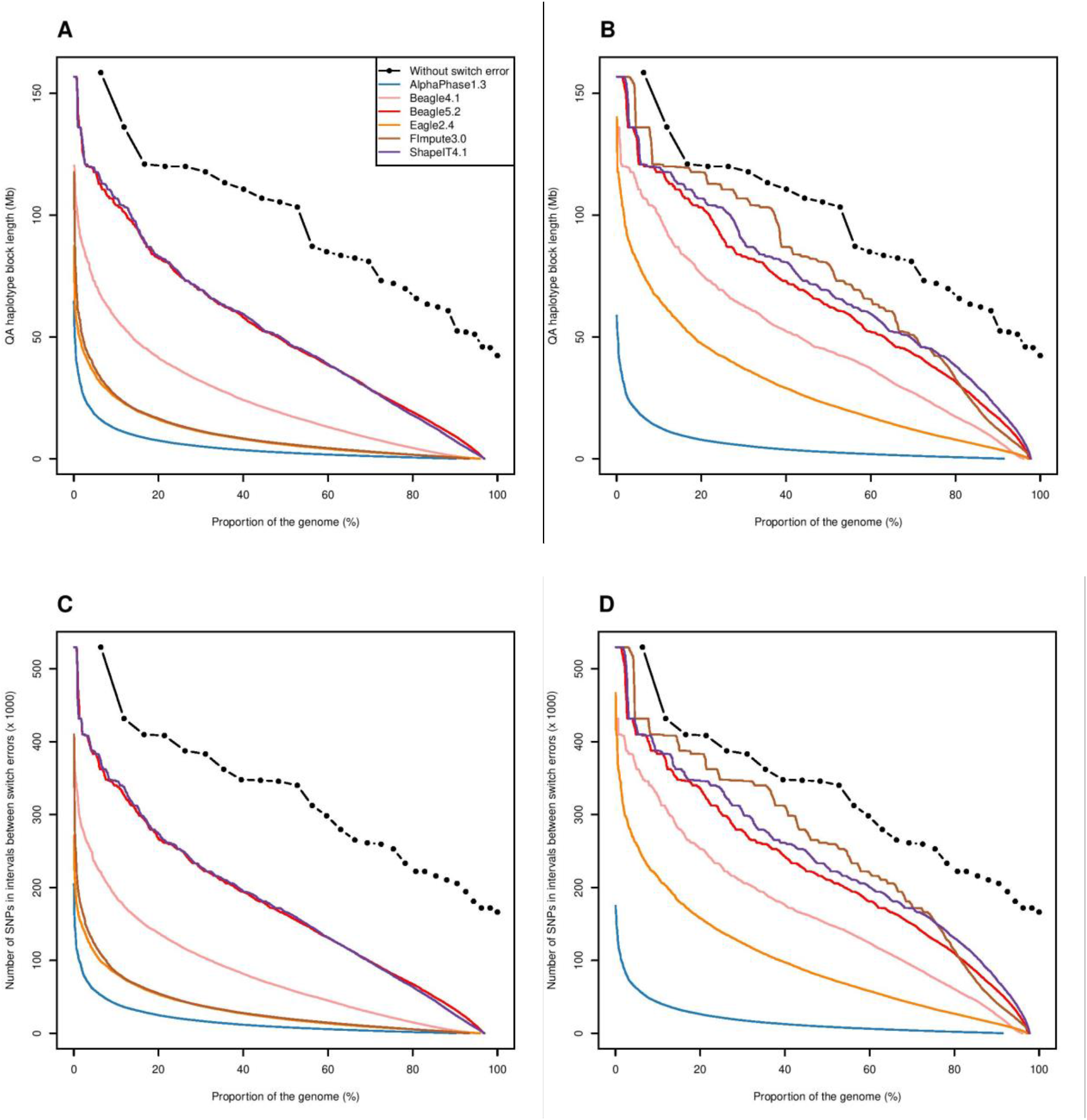
Haplotype block length metrics in both scenarios. (A, B) Quality adjusted (QA) haplotype block lengths (Mb), and (C, D) number of SNPs in correctly phased blocks (x 1000), obtained with AlphaPhase1.3, Beagle4.1, Beagle5.2, Eagle2.4, FImpute3.0 and ShapeIT4.1, computed for the 98 validation individuals and plotted as a function of the proportion of the genome, in scenario 1 (A, C) with only the 98 validation individuals, and in scenario 2 (B, D) with the 264 sequenced individuals. The black curves with dots represent the total length of each chromosome (if they were perfectly phased) as a function of the proportion of the genome they cover.

**Table 2.** Genome percentage included in correctly phased segments longer than different thresholds in both scenarios. Percentage of the genome covering quality adjusted (QA) haplotype blocks of minimal length of respectively 1, 5, 10 and 50 Mb, obtained with AlphaPhase1.3, Beagle4.1, Beagle5.2, Eagle2.4, FImpute3.0 and ShapeIT4.1, computed for the 98 validation individuals in each scenario (scenario 1: using the 98 validation individuals; scenario 2: using the 264 sequenced individuals).

| QA haplotype<br>block length (Mb) | 1 |  | 5 |  | 10 |  | 50 |  |
| --- | --- | --- | --- | --- | --- | --- | --- | --- |
| Scenario | 1 | 2 | 1 | 2 | 1 | 2 | 1 | 2 |
| AlphaPhase1.3 | 71.5 | 73.7 | 30.4 | 32.2 | 13.6 | 14.5 | 0.2 | 0.3 |
| Beagle4.1 | 90.4 | 94.9 | 78.5 | 91.8 | 66.5 | 87.7 | 13.8 | 43.1 |
| Beagle5.2 | 96.3 | 97.2 | 93.6 | 96.1 | 89.3 | 93.9 | 48.1 | 63.2 |
| Eagle2.4 | 86.4 | 96.2 | 54.0 | 87.2 | 33.2 | 74.4 | 1.3 | 18.2 |
| FImpute3.0 | 85.6 | 97.5 | 56.1 | 95.2 | 34.0 | 91.6 | 2.1 | 70.4 |
| ShapeIT4.1 | 96.1 | 97.6 | 92.2 | 96.8 | 87.6 | 95.1 | 48.8 | 69.2 |

In the second scenario including more individuals, mean QA haplotype block lengths (**Table S4** in **Additional file 1**) increased for all methods, reaching 23.3 and 24.1 Mb with Beagle5.2 and ShapeIT4.1, respectively. Interestingly, the improvement is only modest with AlphaPhase1.3 whereas the mean QA haplotype block length increases from 500 kb to 12.5 Mb with FImpute3.0, a 25-fold change. FImpute3.0 is even the best method with respect to QAN50 (79.9 Mb), followed by ShapeIT4.1 (67.1 Mb) and Beagle5.2 (62.7 Mb) (**Table 1**). However, a larger proportion of the genome is included in correctly phased segments longer that 10 Mb with ShapeIT4.1 (95.1%) compared to FImpute3.0 (91.6%) (**Figure 4B,D** and **Table 2**). ShapeIT4.1 performed slightly better than Beagle5.2 at different thresholds (see **Figure 4B,D**). Comparisons of different versions of tested software is in agreement with comparisons made with the first scenario, the differences between Beagle5.0, Beagle5.1 and Beagle5.2 being however smaller (**Figure S2B,D** and **Table S4** in **Additional file 1**).

#### Pairwise SNP phasing accuracy

This metric represents the probability that two SNPs are correctly phased as a function of their distance. The results are reported in **Figure 5** and **Table 3** and are in agreement with observations for other metrics such as QAN50. In the first scenario, these probabilities are above 0.95 and 0.92 at 10 and 100 kb, respectively, with Beagle4.1, Beagle5.2 and ShapeIT4.1 (**Figure 5A**). Other methods presented values below 0.93 and 0.91, respectively. The probabilities dropped rapidly at longer distances, even at 1 Mb (around 0.80 with Beagle4.1 and even below 0.70 for the three less efficient methods). ShapeIT4.1 and Beagle5.2 performed best with probabilities still above 0.90 at 1 Mb, but only 0.78 and 0.66 at 5 and 10 Mb, respectively. At 50 Mb, the probabilities were almost null with all methods and only 0.16 and 0.17 with Beagle5.2 and ShapeIT4.1, respectively. In the second scenario, the probabilities are higher and drop less rapidly, presenting a plateau until a distance of almost 1 Mb (**Figure 5B**). ShapeIT4.1 achieved the highest probabilities, equal to 0.95, 0.87 and 0.77 at 1, 5 and 10 Mb distance, respectively, but FImpute3.0 achieved almost identical results and was even better at very long distance (0.31 at 50 Mb *vs* 0.27 for ShapeIT4.1).

**Figure 5.**
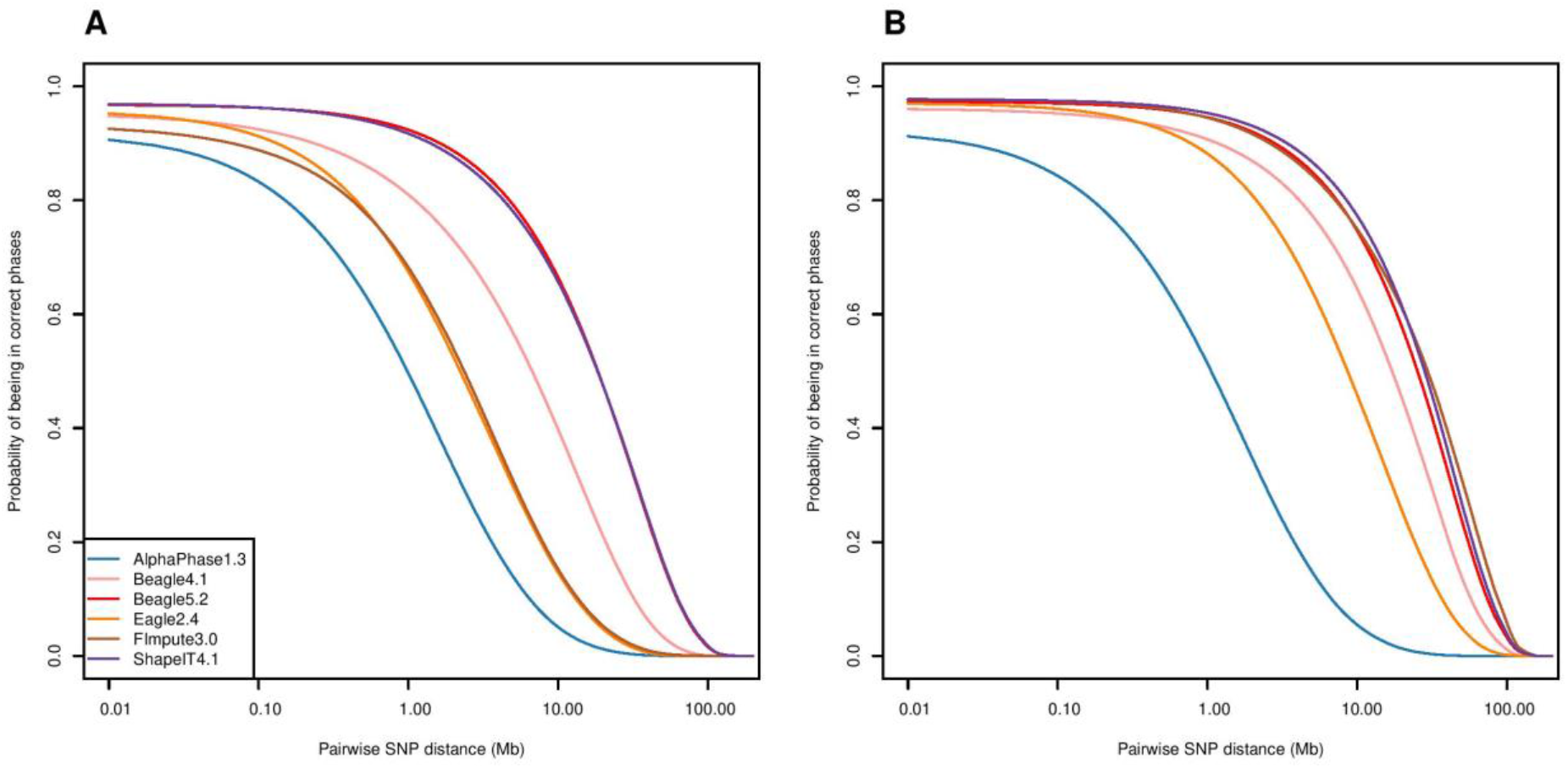
Pairwise SNP phasing accuracy in both scenarios. Probabilities that two SNP are correctly phased as a function of their distance obtained with AlphaPhase1.3, Beagle4.1, Beagle5.2, Eagle2.4, FImpute3.0 and ShapeIT4.1, computed for the 98 validation individuals (A) in scenario 1 with only the 98 validation individuals, and (B) in scenario 2 with the 264 sequenced individuals.

**Table 3.** Pairwise SNP phasing accuracy at different distances in both scenarios. Pairwise SNP phasing accuracy at distances of respectively 0.01, 0.1, 1, 2, 5, 10 and 50 Mb, obtained with AlphaPhase1.3, Beagle4.1, Beagle5.2, Eagle2.4, FImpute3.0 and ShapeIT4.1, computed for the 98 validation individuals in each scenario (scenario 1: using the 98 validation individuals; scenario 2: using the 264 sequenced individuals).

| Pairwise SNP distance (Mb) | 0.01 |  | 0.10 |  | 1 |  | 2 |  | 5 |  | 10 |  | 50 |  |
| --- | --- | --- | --- | --- | --- | --- | --- | --- | --- | --- | --- | --- | --- | --- |
| Scenario | 1 | 2 | 1 | 2 | 1 | 2 | 1 | 2 | 1 | 2 | 1 | 2 | 1 | 2 |
| AlphaPhase1.3 | 0.91 | 0.91 | 0.83 | 0.84 | 0.50 | 0.51 | 0.33 | 0.35 | 0.14 | 0.15 | 0.05 | 0.05 | 0.00 | 0.00 |
| Beagle4.1 | 0.95 | 0.96 | 0.92 | 0.95 | 0.81 | 0.91 | 0.73 | 0.87 | 0.57 | 0.77 | 0.40 | 0.64 | 0.03 | 0.14 |
| Beagle5.2 | 0.97 | 0.97 | 0.96 | 0.97 | 0.92 | 0.95 | 0.89 | 0.92 | 0.79 | 0.85 | 0.66 | 0.75 | 0.16 | 0.24 |
| Eagle2.4 | 0.95 | 0.97 | 0.91 | 0.96 | 0.67 | 0.88 | 0.52 | 0.81 | 0.30 | 0.64 | 0.15 | 0.46 | 0.00 | 0.05 |
| FImpute3.0 | 0.93 | 0.98 | 0.89 | 0.97 | 0.68 | 0.95 | 0.54 | 0.92 | 0.31 | 0.84 | 0.15 | 0.75 | 0.00 | 0.31 |
| ShapeIT4.1 | 0.97 | 0.98 | 0.96 | 0.97 | 0.92 | 0.95 | 0.88 | 0.93 | 0.78 | 0.87 | 0.66 | 0.77 | 0.17 | 0.27 |

## Discussion

We herein compared accuracy of population-based phasing tools in a whole-genome sequenced cattle pedigree. To be able to measure the accuracy, it was essential to apply certain procedures to our whole-genome sequencing data. Indeed, after filtering variants based on a standard variant quality score recalibration procedure, the number of CO in our pedigree was still highly inflated, suggesting that these pedigree-based haplotypes contain many errors. We had to apply further filters to our data to remove additional low-quality markers or small genomic regions incorrectly mapped in the reference genome build. Refining genotype calling with Beagle4.1 [39] had a major impact, stressing the importance of such a procedure. Our final data presented still a few more CO than those obtained at lower density with a high-confidence map. This could be due to the CO missed at lower density or to the errors that remain in the sequence data. This would represent a maximum of 5 incorrect CO on average per meiosis, and these errors could result from a phasing error in the parent or in the offspring haplotypes (each of these haplotypes would thus have less than 5 errors). We could apply additional filters to further reduce the number of errors. For instance, we could remove SNPs located in copy number variants since they would generate spurious CO as genotype calling is more difficult at these positions [41]. Similarly, heterozygous genotypes in the middle of long homozygous-by-descent segments [42] are also probably errors and would generate incorrect CO. Nevertheless, our results illustrate that many errors are still present in whole-genome sequencing data, and that stringent filtering is required. The presence of these low-quality variants would not be a problem in genome-wide association studies because associations are tested independently for each SNP, and genomic prediction methods might be robust to this problem. However, for applications relying on haplotypes and their length, stringent filtering is essential, in particular when long correctly phased segments are required.

The quality of our final data set was high enough to evaluate the phasing methods, since we expect only a few errors (from 0 to 5) in our reference haplotypes. In our best scenario, the accuracy of the best methods was impressive with a median of 50 SEC per individual, corresponding to approximately only two SEC per chromosome. In the case of point switches, a phasing error at a single marker (or a small segment) that would cause two consecutive SEC (see example in **Figure 2**) would represent only one such punctual error per chromosome. Some individuals presented only 2 SEC for their entire genome, and chromosomes were frequently phased without errors. These results are also confirmed with the metrics related to length of correctly phased segments. On average, ShapeIT4.1 had only one switch error every 40,000 informative markers. When only one hundred individuals were simultaneously phased, there were around 10 switches on average per chromosome with Beagle5.2 (one switch error every 10,000 SNPs). These are nevertheless excellent results given the small sample size. These are indeed better results than those reported in human’s populations. For instance, Delaneau *et al.* [35] obtained a SER above 0.5% with ShapeIT4 and Beagle5 and with a reference panel of 20,000 individuals (at lower marker density). Loh *et al.* [33] obtained also higher SER using ShapeIT2 and Eagle2 whereas Choi *et al.* [23] estimated that the SER ranged from 0.8 to 1.5% for Eagle2, ShapeIT2 and Beagle4 for a reference ‘Genome-In-A-Bottle’ whole-genome phased individual, and using a reference panel of 2,500 individuals. Similarly, Song *et al.* [13] reported higher SER, above 2%, in human populations phased with ShapeIT2. This higher accuracy in our cattle data set might be related to the lower effective population size, the higher relatedness and LD levels, particularly at long distance. We previously observed that population-based methods such as Beagle are very effective at indirectly exploiting the familial information through the presence of long-shared haplotypes (see also [24]). Similarly, methods such as AlphaPhase [29] or FImpute [30] can identify parents or surrogate parents without pedigree information. This was confirmed in the present study as increasing the sample size and including sequenced relatives clearly improved the accuracy, in particular for FImpute3.0, although the pedigree information was not explicitly used.

In our study, ShapeIT4.1 and Beagle5.2 performed best for almost all evaluated metrics and for both scenarios. Their relative ranking varied however according to the metric and the scenario. Beagle5.2 achieved the best results mainly in the first scenario whereas ShapeIT4.1 was often 20 the most accurate in the second scenario. When the parents were included in the data, FImpute3.0 accurately phased extremely long segments and the estimated SEC was as low as 2 for some individuals. Nevertheless, when the parents were not included in the sample, the accuracy of FImpute3.0 decreased although some full-sibs were present in the sample (**Figure S3** in **Additional file 1**). Phasing accuracy varies across different versions of a software. In our study, we observed that phasing accuracy improved as expected with newer versions of the software. As a result, comparisons of different methods might vary through time, according to the compared versions. For instance, until recently ShapeIT4.1 was in competition with Beagle5.1. In our comparisons, ShapeIT4.1 was most often better although Beagle5.1 performed extremely well. However, Beagle5.2, the new release, performed as well as ShapeIT4.1 (see above). Phasing accuracy will also change according to different elements such as marker density, level of relatedness and size of the population [24], and this might impact the ranking of the methods. For instance, in Choi *et al.* [23], Eagle2 performed better than ShapeIT2 and Beagle4 on human data. In a Holstein dairy cattle population genotyped with medium to high density genotyping arrays, Miar *et al.* [43] compared Beagle4.1, ShapeIT2 and FImpute based on SER. They estimated that Beagle4.1 was the most accurate whereas ShapeIT2 resulted in higher SER. However, their sample was much larger and more information from relatives was thus available. Consistently with our study, when one or two parents of the validation animals were added to the phased sample, FImpute became more accurate than Beagle4.1. Using simulated data mimicking a brown layer population, Frioni *et al.* [44] observed that haplotypes phased with Beagle4.1 had lower SEC than those obtained with FImpute when parents were not included. As in our study, inclusion of parents in the phased sample increased phasing accuracy for FImpute. Fewer comparisons in livestock species are available for ShapeIT4.1 or Beagle5.0 as these programs are more recent. In summary, our data set represent a typical example of reference panel containing 100 to 200 whole-genome sequenced individuals in a livestock species with high levels of relatedness. In those conditions, ShapeIT4.1 and Beagle5.2 performed particularly well. In addition, we observed that accuracy of ShapeIT4.1 could be further improved by optimizing parameters (by increasing the value of the `--pbwt-depth` parameter for instance).

The choice of the phasing method might nevertheless depend on the availability of other options. For instance, Beagle4.0, FImpute3.0 and AlphaPhase1.3 can exploit the pedigree information, which might increase their phasing accuracy in certain conditions. We previously proposed a two-step procedure in which haplotypes are first obtained based on familial information and unphased markers are subsequently phased by a LD-based approach [26]. Such an approach is also possible with Beagle4.1 or ShapeIT4.1, that will preserve pre-phasing information present in the VCF file. Phasing information coming from marker alleles present on the same sequenced reads can also be integrated with such an approach.

Finally, the importance of phasing accuracy will depend on the applications in which the haplotypes are used. For many applications, accurate phasing is only required at short range. For haplotype-based association studies, short 100-kb haplotypes would capture interactions among tightly-linked *loci*. We previously observed that improved long-range phasing accuracy did not result in higher imputation accuracy in a livestock population [45]. The presence of a few switch errors would not necessarily be a problem in haplotype-based GWAS or genomic selection, or in some QTL mapping approaches, as long as correctly phased segments are long enough to infer the IBD relationships around the tested position. For such applications, most of the tested methods would provide sufficient accuracy. The phasing accuracy will be more important in applications in which the length of shared haplotypes is used to estimate age of alleles [17] or age to a common ancestor, to identify signatures of selection [15, 16], to determine relatedness between individuals based on the distribution of length of shared IBD segments [4]. This accuracy will also be essential in studies on meiotic recombination based on the identification of CO in genotyped or sequenced pedigrees [20, 46].

## Methods

### Sequencing data

The whole-genome sequence data used in the present work was obtained from 264 Holstein-Friesian individuals from the DAMONA pedigree designed to study germline mutation in cattle [47] and previously used and described [48, 49]. The individuals were sequenced at high coverage (mean coverage: 25.8X, ranging from 15.2X to 47.1X), and the data included 98 sequenced trios (**Figure S3** in **Additional file 1**). Whole genome Illumina Nextera PCR free libraries (550 bp insert size) were sequenced on an Illumina HiSeq 2000 with a paired-end protocol (2×100 bp).

The sequencing data was re-aligned on the new ARS-UCD1.2 (BosTau9) bovine genome assembly [37] using the Burrows-Wheeler Aligner MEM algorithm (v0.7.5a) [50]. The SAM files were converted into BAM files with SAMtools (v1.9) [51]. The BAM files were sorted using Sambamba (v0.6.6) [52]. PCR duplicates were marked with the MarkDuplicates option of picard-tools (v2.7.1) [53]. The BAM files were then recalibrated using the BaseRecalibrator procedure of GATK (v4.1.7.0) [54–56], using the VCF provided by the 1000 Bull Genome project (http://www.1000bullgenomes.com/) as known polymorphic sites database. Individual GVCF files were obtained with HaplotypeCaller (GATK4) and were subsequently merged in a GenomicsDB (with GenomicsDBImport, GATK4) to perform joint genotyping with GenotypeGVCFs (GATK4). Variants from the resulting VCF file were then recalibrated using VariantRecalibrator (GATK4) by applying two thresholds (99.9 and 97.5) and using 1.2M SNPs extracted from commercial chips [57] as truth and training sets, and 138M SNPs provided by the 1000 Bull Genome project as known set.

### Assessing the quality of whole-genome sequence genotype data

#### Strategy

To evaluate the quality of whole-genome sequence data in terms of genotyping error rates and of physical marker order, we compared the number of CO identified in our pedigree with the sequence data to the number of CO identified in our pedigree with a lower density high-confidence map validated on a much larger population genotyped with the Illumina BovineSNP50 BeadChip (Illumina Inc., San Diego, CA). The CO are identified in sequenced parent-offspring pairs, also referred to as proband and gamete. For the comparisons, we relied only on the most accurate CO counts, obtained when at least one parent of the proband was also sequenced (160 gametes out of 279: 104 females and 56 males probands). Although a few CO might be missed or incorrectly identified with this reference map, much higher counts of CO with the sequence data would indicate presence of genotyping and map errors.

#### Validation of the reference 50K map

To obtain a high-confidence reference 50K marker map, we used the genotype data from Kadri *et al.* [48], available for 115,967 individuals and 30,349 SNPs and followed their approach relying on the map confidence score (MCS) implemented in LINKPHASE3 [36] to identify eventually misplaced markers that were subsequently removed. Physical positions of the markers in the ARS-UCD1.2 bovine genome assembly were obtained from https://www.animalgenome.org/repository/cattle/UMC_bovine_coordinates. Among the 30,331 SNPs from this high-confidence reference map, 29,375 SNPs were present in our whole-genome sequence data. The resulting high-confidence reference 50K marker map is provided in **Additional file 2**.

#### Evaluated data processing steps

##### Variant quality score recalibration (VQSR)

By applying the VariantRecalibrator procedure (GATK, see above) on the genotype calls, we kept 15,327,429 and 11,030,905 SNPs with the 99.9 and 97.5 thresholds, respectively. These variants correspond to 99.9 and 97.5% of the total truth sites, 95.1 and 76.4% of the total known SNPs, and 60.1% and 16.0% of the total novel SNPs, respectively.

##### Variant selection based on genetic rules

To further enrich our data set in high-quality variants, we selected variants behaving as true SNPs based on Hardy-Weinberg equilibrium test (p > 0.05), with expected genotype frequencies in offspring from heterozygous parents (based on a χ² test, p > 0.05), and not presenting more than one Mendelian inconsistency in parent-offspring pairs or trios (e.g., opposite homozygotes) for the entire pedigree. We kept only variants for which the probability to observe no Mendelian inconsistencies by chance was lower than 1e-12. Finally, we also discarded uninformative markers with low minor allele frequency (< 0.01). Applications of these rules resulted in the selection of 8,435,899 informative variants behaving as true SNPs.

##### Improving genotyping-calling

The accuracy of genotyping-calling for these 8,435,899 variants was then improved with the LD-based approach implemented in Beagle4.1 [39].

##### Exclusion of suspicious genomic regions

Several filters were applied to exclude genomic regions of putatively lower quality. First, we removed the last 10-kb segments at chromosome ends. Second, we excluded high coverage regions that we identified within each individual using the approach described in the LUMPY framework [58]. We considered as high coverage regions the genomic regions presenting 6 times higher coverage than the individual average whole-genome coverage. Finally, we discarded genomic regions associated with putative errors in the genome assembly. To that end we used tools available in LINKPHASE3 [36] and an approach described in more details in Kadri *et al.* [41]. We first identified small genomic regions flanked by peaks of high recombination rate (> 0.05). For these genomic regions, we compared within-family segregation patterns (inheritance vectors as described in Druet and Georges [36]) to those obtained with the high-confidence 50K map in the flanking regions, and removed those with a r² < 0.90. We subsequently removed SNPs with a low MCS (< 0.99) as well as chromosome extremities when they presented inflated RR with flanking regions. The procedure was repeated until no further evidence for map errors was visible. In total, we excluded 18,220 SNPs associated with these regions (**Additional file 3**).

##### Final editions

We finally set to missing 38,537 incompatible genotypes in parent-offspring pairs (opposite homozygous), eventually introduced after running Beagle4.1. We also removed SNPs that were monomorphic at this stage and those with a missing genotyping rate above 5% in the final pedigree. The final number of SNPs was 8,417,679 (complete list provided in **Additional file 4**).

### Phasing methods

We herein compared the accuracy from different phasing software: AlphaPhase v1.1 [29] and v1.3 [59], ShapeIT v2.r904 [31] and v4.1.3 [35], Beagle v3.3.2, v4.0, v4.1, v5.0, v5.1, and v5.2 [7, 60], Eagle v2.4.1 [33], and FImpute v3.0 [30]. Most of these methods are population-based methods. Indeed, Eagle and ShapeIT are hidden Markov models (HMM) modeling target haplotypes as mosaic of reference haplotypes, similarly to the Li and Stephens model [40]. The earlier versions of Beagle (v3.3.2, v4.0 and v4.1 [7]) condense all the observed haplotypes into a directed acyclic graph and the resulting model is a variable length Markov chain that can also be viewed as a HMM. The newer versions of Beagle (v5.0, v5.1 and v5.2 [60]) are based on the Li and Stephens HMM [40]. AlphaPhase and FImpute are both heuristic methods performing haplotype matching in fixed-length windows, and that exploit also the pedigree information when possible. More detailed descriptions of these models are available in the original papers. For each of the tools, we used the default parameters.

Comparisons were first realized with the most recent version of different software, including AlphaPhase1.3, ShapeIT4.1, Beagle4.1, Beagle5.2, Eagle2.4 and FImpute3.0. We included Beagle4.1 because the method is very different from Beagle5.2. Then, we compared different versions of some software to evaluate the benefit of different updates and to obtain metrics for older versions that have been compared in previous studies, or that are sometimes still in use. This allows also to evaluate whether it is worth re-phasing a data set after the release of a new version. FImpute3.0 and Eagle2.4 were not included in this comparison as we only tested one version for these software.

### Evaluation of haplotype phasing quality

#### Gold standard for haplotype evaluation

In real data sets, true haplotypes remain generally unknown. Therefore, we rely on a family-based strategy that is possible thanks to the availability of a sequenced pedigree [24, 61]. In that case, child’s haplotypes can be resolved using Mendelian segregation rules when at least one of their parents is also sequenced. In the absence of genotyping or marker order errors, this results in exact haplotypes although some positions might remain unphased. To minimize those errors in our data set, we applied the different procedures described above. The family-based haplotypes can then be used to assess the quality of haplotypes obtained using population-based phasing methods. In our sequenced pedigree, we had 98 sequenced offspring with both parents sequenced (trios). We phased these trios using Mendelian segregation rules with LINKPHASE3 [36] and used them subsequently as the gold standard (referred to as “true haplotypes”). In the first tested scenario, the sample of 98 offspring (validation individuals) was phased with each method. In a second scenario, all the 264 sequenced individuals were included in the phasing step, including the sequenced parents but the pedigree information was ignored. This second scenario allowed to study the impact of increasing the sample size and including more related individuals.

#### Metrics for phasing performance

##### Phasing yield

This metric defined as the percentage of phased single nucleotide variants, and representing the completeness of phased haplotypes, is sometimes used in evaluation of phasing quality (e.g., [23]). However, all the evaluated population-based methods achieve 100% phasing yield and this metric is therefore only relevant for AlphaPhase and FImpute.

##### Switch error counts and rates

For each evaluated phasing method, we compared within individual the combinations of alleles at each pair of consecutive informative sites (i.e. heterozygous and phased in the true haplotype) in the inferred haplotypes with the combination of alleles present in the true haplotypes as illustrated in **Figure 2**. Each discrepancy is called a switch error and the total number of switch errors per individual is the switch error count (SEC). Since this number depends on the number of informative SNPs, that represents also the number of opportunities for switch errors, we also defined the switch error rate (SER) as the SEC divided by the number of informative markers (e.g., [23]).

##### Length of correctly phased haplotype blocks

In order to characterize the distance between successive switch errors (distribution along the genome), we used the quality adjusted (QA) haplotype block length and the QAN50 metrics described in Duitama *et al.* [62]. Briefly, the QA haplotype block length is defined as the length of a segment between two successive switch errors (**Figure 2**) multiplied by the proportion of phased SNPs. The QAN50 is defined as the longest QA length such that 50% of all the informative SNPs are located within haplotype blocks with a QA length larger than QAN50 [4]. In other words, this means that 50% of the informative SNPs are contained in haplotype blocks of at least QAN50. Note that the quality adjustment was only relevant for methods with a phasing yield below 100%. For other methods, these metrics represent raw block lengths.

##### Pairwise SNP phasing accuracy

Finally, we computed the probabilities for two SNPs to be correctly phased as a function of the distance between SNPs pairs, providing a complementary measure to previous metrics [23].

## Declarations

### Ethics approval and consent to participate

Not applicable.

### Consent for publication

Not applicable.

### Availability of data and materials

All relevant data supporting the conclusions of this article are included in the article and its supplementary files. The datasets analyzed during the current study are not yet publicly available because they will be released after the publication of another study, but are available from the corresponding author on reasonable request.

### Authors’ contributions

LK, WC, GCMM and MG provided data and resources. CO, NK and TD performed the experiments, analyzed and interpreted the data. CO and TD wrote the manuscript. All authors read and approved the final manuscript.

## Acknowledgements

We thank Erik Mullaart and CRV (Arnhem, The Netherlands) for providing the samples. Tom Druet is Senior Research Associate from the Fonds de la Recherche Scientifique - FNRS (F.R.S.-FNRS). Computational resources have been provided by the *Consortium des Équipements de Calcul Intensif* (CÉCI), funded by the Fonds de la Recherche Scientifique - FNRS (F.R.S.-FNRS) under Grant No. 2.5020.11 and by the Walloon Region. The authors also acknowledge use of the GIGA high performance computing cluster for conducting the study reported in this paper.

## Supplementary Information

### Additional file 1

Supplementary Figures (S1-S3) and Tables (S1-S4). **Figure S1.** Switch error rates in earlier versions of evaluated software in both scenarios. **Figure S2.** Haplotype block length metrics in earlier versions of evaluated software in both scenarios. **Figure S3.** Pedigree tree of the 264 Holstein-Friesian cattle used in this study. **Table S1.** Number of SNPs per chromosome in both scenarios. **Table S2**. Summary statistics of the switch error counts in both scenarios. **Table S3**. Summary statistics of the switch error rates in both scenarios. **Table S4.** Summary statistics of the quality adjusted haplotype block lengths in both scenarios.

### Additional file 2

**High-confidence reference 50K marker map.** Each line of the table corresponds to an interval between two successive SNPs with the following columns: chromosome number / marker name of the first SNP / physical position (bp/10,000,000) of the first SNP / marker name of the second SNP / physical position (bp/10,000,000) of the second SNP / estimated genetic distance (Morgans) in males / estimated genetic distance (Morgans) in females.

### Additional file 3

**Suspicious genomic regions excluded in this study.** Each line of the table corresponds to a genomic region with the following columns: chromosome number / physical start position (bp) / physical end position (bp).

### Additional file 4

**Full list of the sequence SNPs used in this study.** Each line of the table corresponds to a SNP with the following columns: chromosome number / physical position (bp).

## List of abbreviations

CO: Cross-Over
IBD: Identical-By-Descent
LD: Linkage Disequilibrium
MCS: Map Confidence Score
QA: Quality Adjusted
QTL: Quantitative-Trait Locus
SEC: Switch Error Count
SER: Switch Error Rate
VQSR: Variant Quality Score Recalibration

