## Additional file 1 for "Benchmarking phasing software with a whole-genome sequenced cattle pedigree"

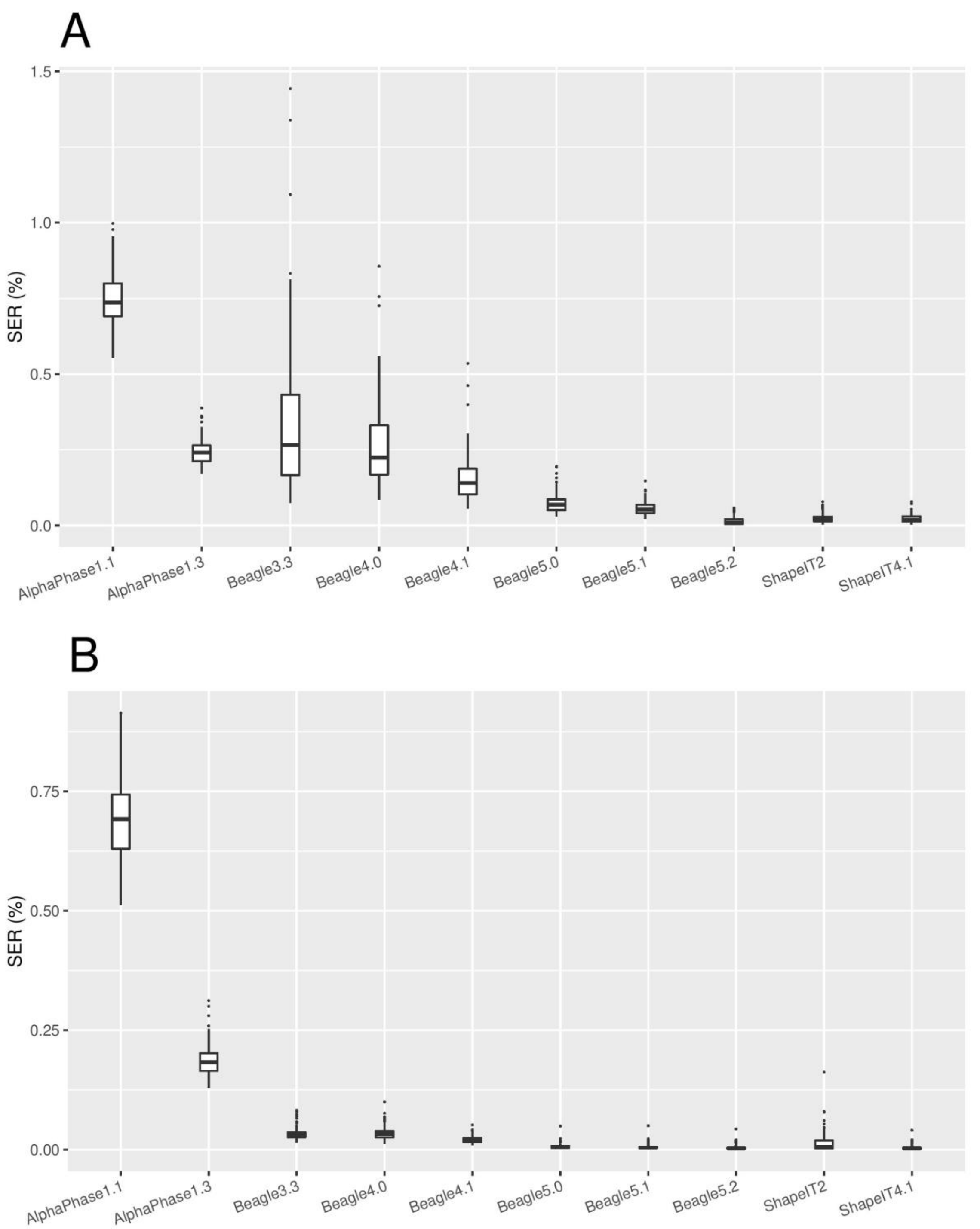


**Figure S1. Switch error rates in earlier versions of evaluated software in both scenarios.** *Boxplots of switch error rates (SER, %) obtained with AlphaPhase1.1, AlphaPhase1.3, Beagle3.3, Beagle4.0, Beagle4.1, Beagle5.0, Beagle5.1, Beagle5.2, ShapeIT2 and ShapeIT4.1, computed for the 98 validation individuals (A) in scenario 1 with only the 98 validation individuals, and (B) in scenario 2 with the 264 sequenced individuals.*

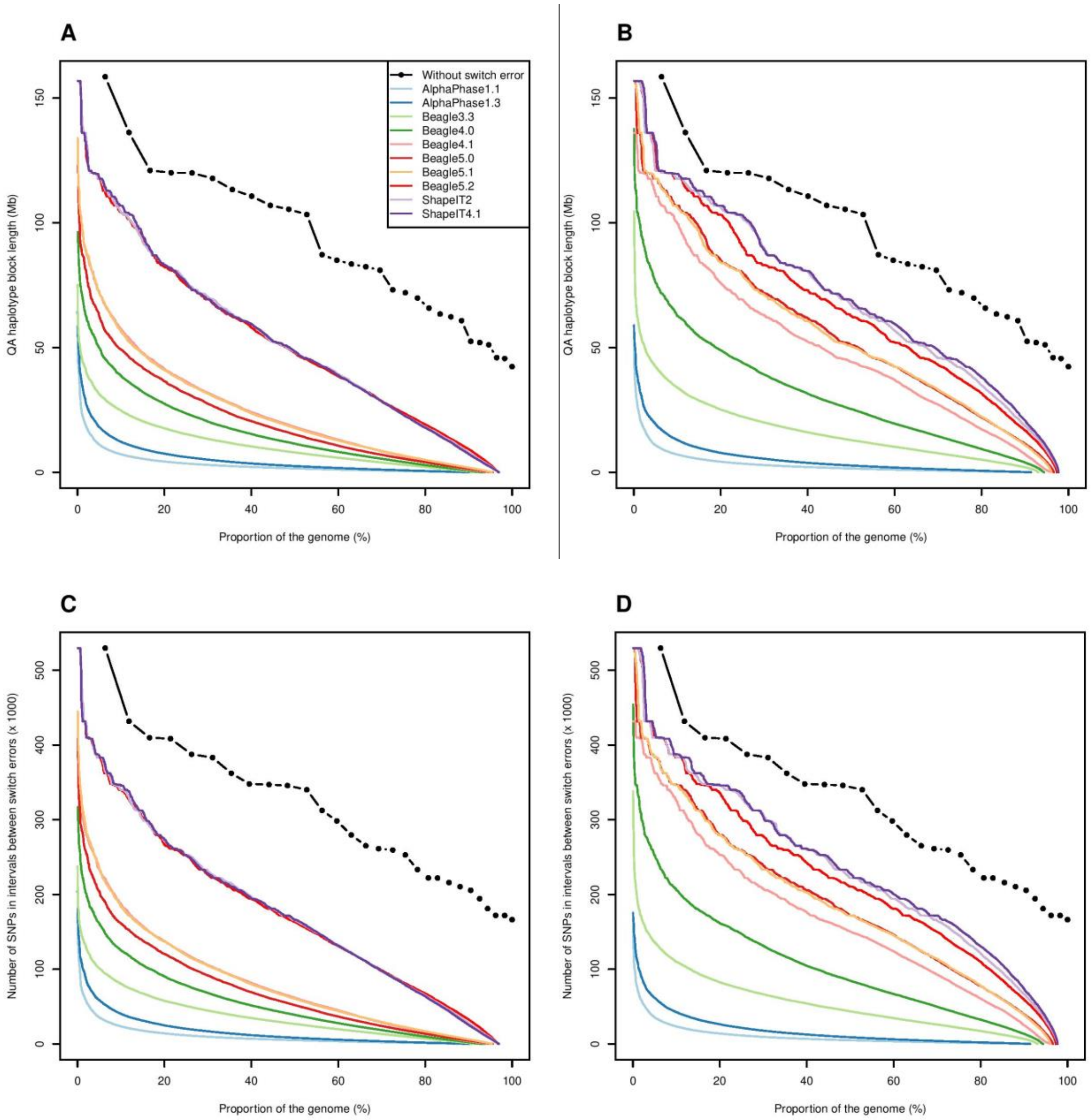


**Figure S2. Haplotype block length metrics in earlier versions of evaluated software in both scenarios.** *(A, B) Quality adjusted (QA) haplotype block lengths (Mb), and (C, D) number of SNPs in correctly phased blocks (x 1000), obtained with AlphaPhase1.1, AlphaPhase1.3, Beagle3.3, Beagle4.0, Beagle4.1, Beagle5.0, Beagle5.1, Beagle5.2, ShapeIT2 and ShapeIT4.1, computed for the 98 validation individuals and plotted as a function of the proportion of the genome, in scenario 1 (A, C) with only the 98 validation individuals, and in scenario 2 (B, D) with the 264 sequenced individuals. The black curves with dots represent the total length of each chromosome (if they were perfectly phased) as a function of the proportion of the genome they cover.*

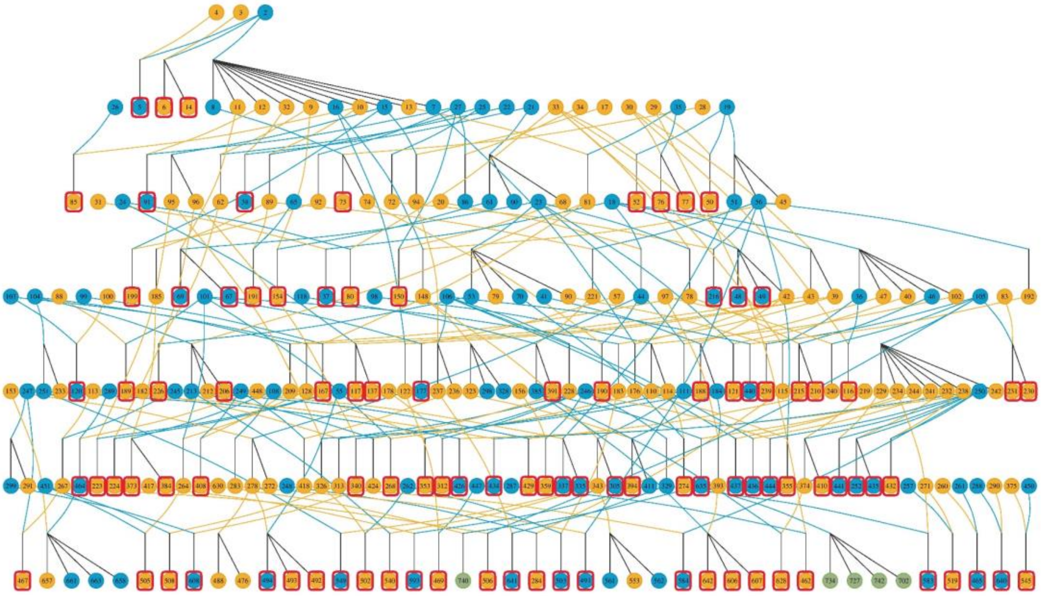

**Figure S3. Pedigree tree of the 264 Holstein-Friesian cattle used in this study**. *Individuals in blue are males, the ones in orange are females, and the ones surrounded in red are the 98 validation individuals (offspring from trios, that is with both parent sequenced) used to compare the algorithms.*

| Chromosome | Number of SNPs after applying filtering procedures (see *Methods*) | Number of SNPs after removing monomorphic SNPs and SNPs with a missing genotyping rate below 5% | | Mean number of heterozygous SNPs across the 98 offspring | |
|---|---|---|---|---|---|
| | | Scenario 1 | Scenario 2 | Scenario 1 | Scenario 2 |
| 1 | 529,626 | 529,582 | 529,600 | 121,003 | 121,002 |
| 2 | 431,779 | 431,702 | 431,719 | 102,524 | 102,524 |
| 3 | 387,566 | 387,531 | 387,547 | 88,918 | 88,918 |
| 4 | 408,467 | 408,433 | 408,446 | 96,956 | 96,956 |
| 5 | 383,157 | 383,127 | 383,146 | 89,821 | 89,822 |
| 6 | 409,796 | 409,756 | 409,785 | 98,721 | 98,721 |
| 7 | 340,141 | 340,070 | 340,126 | 79,742 | 79,742 |
| 8 | 346,970 | 346,941 | 346,964 | 79,739 | 79,739 |
| 9 | 345,865 | 345,644 | 345,847 | 79,126 | 79,126 |
| 10 | 347,854 | 347,839 | 347,844 | 79,916 | 79,915 |
| 11 | 362,191 | 362,170 | 362,179 | 84,009 | 84,009 |
| 12 | 312,506 | 312,477 | 312,491 | 73,220 | 73,221 |
| 13 | 265,193 | 265,167 | 265,183 | 61,133 | 61,133 |
| 14 | 279,542 | 279,528 | 279,534 | 64,147 | 64,147 |
| 15 | 298,393 | 298,364 | 298,372 | 69,585 | 69,586 |
| 16 | 259,508 | 259,454 | 259,497 | 61,050 | 61,050 |
| 17 | 261,305 | 261,292 | 261,300 | 62,710 | 62,710 |
| 18 | 215,908 | 215,900 | 215,904 | 50,969 | 50,969 |
| 19 | 210,452 | 210,447 | 210,450 | 49,474 | 49,474 |
| 20 | 252,881 | 252,860 | 252,875 | 59,498 | 59,498 |
| 21 | 233,326 | 233,248 | 233,308 | 54,239 | 54,239 |
| 22 | 205,619 | 205,611 | 205,617 | 48,671 | 48,671 |
| 23 | 222,026 | 222,000 | 222,018 | 51,325 | 51,326 |
| 24 | 222,142 | 222,129 | 222,134 | 51,827 | 51,827 |
| 25 | 166,292 | 166,279 | 166,286 | 39,031 | 39,031 |
| 26 | 171,810 | 171,803 | 171,799 | 38,886 | 38,884 |
| 27 | 181,041 | 181,030 | 181,034 | 43,365 | 43,365 |
| 28 | 172,002 | 171,992 | 171,997 | 38,908 | 38,908 |
| 29 | 194,321 | 194,306 | 194,318 | 45,708 | 45,709 |
| Total | 8,417,679 | 8,416,682 | 8,417,320 | 1,964,220 | 1,964,220 |

**Table S1. Number of SNPs per chromosome in both scenarios.** *Total number of whole-genome sequence SNPs and mean numbers of informative SNPs for each chromosome in each scenario (scenario 1: using the 98 validation individuals; scenario 2: using the 264 sequenced individuals).*

| Metric | Min. SEC | | Mean SEC | | Median SEC | | Max. SEC | |
|---|---|---|---|---|---|---|---|---|
| Scenario | 1 | 2 | 1 | 2 | 1 | 2 | 1 | 2 |
| AlphaPhase1.1 | 10,790 | 10,338 | 14,767 | 13,763 | 14,341 | 13,439 | 19,930 | 18,982 |
| AlphaPhase1.3 | 3,339 | 2,635 | 4,796 | 3,698 | 4,612 | 3,548 | 7,675 | 6,169 |
| Beagle3.3 | 1,452 | 289 | 6,595 | 643 | 5,209 | 600 | 28,854 | 1,596 |
| Beagle4.0 | 1,655 | 241 | 5,121 | 657 | 4,435 | 648 | 16,925 | 1,940 |
| Beagle4.1 | 1,032 | 196 | 3,113 | 399 | 2,769 | 359 | 10,578 | 999 |
| Beagle5.0 | 595 | 30 | 1,491 | 128 | 1,339 | 98 | 3,960 | 949 |
| Beagle5.1 | 455 | 26 | 1,107 | 108 | 1,024 | 81 | 2,913 | 966 |
| Beagle5.2 | 35 | 13 | 293 | 75 | 191 | 55 | 1,136 | 834 |
| Eagle2.4 | 2,799 | 325 | 5,036 | 894 | 4,901 | 882 | 9,031 | 1,535 |
| FImpute3.0 | 3,012 | 2 | 4,603 | 165 | 4,477 | 115 | 7,616 | 598 |
| ShapeIT2 | 64 | 8 | 459 | 307 | 377 | 110 | 1,551 | 3,277 |
| ShapeIT4.1 | 64 | 12 | 449 | 72 | 386 | 50 | 1,561 | 785 |

**Table S2. Summary statistics of the switch error counts in both scenarios.** *Minimum, mean, median and maximum switch error counts (SEC), obtained with all the evaluated LD-based phasing algorithms, computed for the 98 validation individuals in each scenario (scenario 1: using the 98 validation individuals; scenario 2: using the 264 sequenced individuals).*

| Metric | Min. SER (%) | | Mean SER (%) | | Median SER (%) | | Max. SER (%) | |
|---|---|---|---|---|---|---|---|---|
| Scenario | 1 | 2 | 1 | 2 | 1 | 2 | 1 | 2 |
| AlphaPhase1.1 | 0.5547 | 0.5120 | 0.7510 | 0.7000 | 0.7369 | 0.6919 | 0.9977 | 0.9139 |
| AlphaPhase1.3 | 0.1716 | 0.1293 | 0.2440 | 0.1882 | 0.2412 | 0.1834 | 0.3884 | 0.3122 |
| Beagle3.3 | 0.0746 | 0.0145 | 0.3366 | 0.0328 | 0.2657 | 0.0300 | 1.4426 | 0.0825 |
| Beagle4.0 | 0.0851 | 0.0121 | 0.2609 | 0.0336 | 0.2240 | 0.0322 | 0.8564 | 0.1003 |
| Beagle4.1 | 0.0556 | 0.0091 | 0.1583 | 0.0204 | 0.1404 | 0.0183 | 0.5353 | 0.0517 |
| Beagle5.0 | 0.0306 | 0.0015 | 0.0756 | 0.0066 | 0.0688 | 0.0050 | 0.1954 | 0.0491 |
| Beagle5.1 | 0.0227 | 0.0013 | 0.0564 | 0.0056 | 0.0523 | 0.0041 | 0.1474 | 0.0500 |
| Beagle5.2 | 0.0018 | 0.0007 | 0.0150 | 0.0039 | 0.0093 | 0.0027 | 0.0575 | 0.0431 |
| Eagle2.4 | 0.1439 | 0.0167 | 0.2561 | 0.0455 | 0.2485 | 0.0451 | 0.4570 | 0.0753 |
| FImpute3.0 | 0.1371 | 0.0001 | 0.2343 | 0.0085 | 0.2256 | 0.0059 | 0.3854 | 0.0316 |
| ShapeIT2 | 0.0032 | 0.0004 | 0.0234 | 0.0157 | 0.0187 | 0.0056 | 0.0785 | 0.1620 |
| ShapeIT4.1 | 0.0032 | 0.0006 | 0.0229 | 0.0037 | 0.0192 | 0.0026 | 0.0790 | 0.0406 |

**Table S3. Summary statistics of the switch error rates in both scenarios.** *Minimum, mean, median and maximum switch error rates (SER, %), obtained with all the evaluated LD-based phasing algorithms, computed for the 98 validation individuals in each scenario (scenario 1: using the 98 validation individuals; scenario 2: using the 264 sequenced individuals).*

| Metric | Min. QA haplotype block length (bp) | | Mean QA haplotype block length (bp) | | Median QA haplotype block length (bp) | | Max. QA haplotype block length (bp) | |
|---|---|---|---|---|---|---|---|---|
| Scenario | 1 | 2 | 1 | 2 | 1 | 2 | 1 | 2 |
| AlphaPhase1.1 | 1 | 1 | 152,648 | 163,711 | 621 | 725 | 58,851,352 | 59,132,672 |
| AlphaPhase1.3 | 1 | 1 | 465,447 | 610,536 | 10,653 | 55,973 | 64,541,591 | 58,753,065 |
| Beagle3.3 | 1 | 1 | 346,928 | 3,446,510 | 166 | 13,201 | 75,174,431 | 104,596,496 |
| Beagle4.0 | 1 | 1 | 453,121 | 3,426,126 | 337 | 1,893 | 96,286,002 | 137,647,941 |
| Beagle4.1 | 1 | 1 | 754,558 | 5,595,676 | 484 | 45,729 | 120,319,442 | 136,047,557 |
| Beagle5.0 | 1 | 1 | 1,564,542 | 15,344,484 | 9,118 | 1,898,160 | 124,880,023 | 156,851,007 |
| Beagle5.1 | 1 | 1 | 2,090,531 | 17,464,973 | 7,924 | 4,271,136 | 133,971,707 | 156,846,286 |
| Beagle5.2 | 1 | 1 | 7,479,255 | 23,313,031 | 12,334 | 6,953,676 | 156,851,371 | 156,851,371 |
| Eagle2.4 | 1 | 1 | 471,275 | 2,618,868 | 172 | 161 | 87,438,003 | 140,190,109 |
| FImpute3.0 | 1 | 1 | 501,289 | 12,561,540 | 583 | 27,490 | 117,669,102 | 156,851,371 |
| ShapeIT2 | 1 | 1 | 4,950,799 | 7,244,068 | 1,456 | 433 | 156,851,371 | 156,851,371 |
| ShapeIT4.1 | 1 | 1 | 5,050,285 | 24,108,523 | 2,134 | 4,738,542 | 156,851,371 | 156,851,371 |

**Table S4. Summary statistics of the quality adjusted haplotype block lengths in both scenarios.** *Minimum, mean, median and maximum quality adjusted (QA) haplotype block lengths, obtained with all the evaluated LD-based phasing algorithms, computed for the 98 validation individuals in each scenario (scenario 1: using the 98 validation individuals; scenario 2: using the 264 sequenced individuals).*
